# PhenoMS-ML: Phenotypic Screening by Mass Spectrometry and Machine Learning

**DOI:** 10.1101/593244

**Authors:** Luuk N. van Oosten, Christian D. Klein

## Abstract

Protein mass fingerprinting by MALDI-TOF MS in combination with machine learning (PhenoMS-ML) permits the identification of response signatures generated in cell cultures upon exposure to well-characterized drugs. PhenoMS-ML is capable to identify and classify the mode of action of unknown antibacterial agents in wild-type *Escherichia coli* and *Staphylococcus aureus*. It allows the sensitive, specific, and high-throughput identification of drug target mechanisms that are difficult to assess by other methods.

## Main

Compound activity data from assays at isolated target proteins play an important role in pharmacology, toxicology and medicinal chemistry, but their translation into systems of higher complexity such as cell cultures (or patients) is frequently difficult (Brown and Wright 2016). This is caused by pharmacokinetic effects, macromolecular crowding effects in the intracellular environment which are absent in a biochemical buffer, or intracellular presence of competing ligands and substrates, such as ATP (Swinney 2014). Numerous important pharmacological targets are difficult, if not impossible, to study in biochemical systems because of their dependency on a specific environment or unusual substrates. This is particularly evident and problematic in the field of antibacterial drug discovery, where we (Bachelier, Mayer et al. 2006, Schiffmann, Neugebauer et al. 2006, Mendgen, Scholz et al. 2010) and many others (Payne, Gwynn et al. 2006) have repeatedly failed to translate potent biochemical inhibitors into antibacterial drug candidates. Undeterred by the efforts put in the identification of novel targets and mode of actions in bacteria, the main target pathways of new and established antibacterial agents remain cell wall synthesis, ribosomal machinery, and nucleic acid processing (Livermore, Blaser et al. 2011). Making things worse, these pathways are notoriously difficult to study in biochemical systems, let alone in high-throughput manner, as would be desirable for compound screenings.

Considering the numerous difficulties involved in setting up individual assay procedures for these important antibacterial targets, whose results would be a limited predictor for actual *in vivo* efficacy, we reasoned that a phenotypic approach to drug screening is highly desirable. Phenotypic antimicrobial testing is typically performed using growth assays (Silver 2011). However, information obtained from such assays is mostly restricted to a binary ‘dead-or-alive’ information, and does not provide any further information about the targets, pathways, or modes of action that are involved. It seems advantageous to employ cell-based phenotypic screening methods that yield more information on the target and mode of action involved (Feng, Mitchison et al. 2009).

A method that addresses this issue is bacterial cytological profiling as described by the Pogliano group (Nonejuie, Burkart et al. 2013), who identified cellular pathways involved in response to antibiotics by means of fluorescence microscopy. Another example of such a method is Raman spectroscopy profiling of bacteria in response to antibiotic induced stress (Athamneh, Alajlouni et al. 2013). However, a common disadvantage of both methods is that the relative amount of antibiotic required to see an effect is relatively high, over 2× to 5× the minimal inhibitory concentration (MIC), making it impossible to identify weakly active compounds in wild-type bacteria.

The present work is based on the hypothesis that mass spectra obtained from wild-type cells under the influence of chemical stressors provide a fine-grained description of the proteomic state of a cell culture. We further reasoned that this specific response to the stressor can be recognized by state-of-the-art machine learning algorithms and further utilized to screen drug candidates. We show here that proteomic fingerprints of cells treated with known antibiotics can be used to characterize other compounds and pinpoint their effect on antibacterial drug targets. Mass spectra of cell cultures were acquired by matrix assisted laser desorption ionization mass spectrometry (MALDI-TOF MS), a method which requires minimal sample preparation, is high-throughput amenable, and has a long track record in the microbiology field (Kostrzewa 2018).

Bacterial cells of *Escherichia coli* (*E. coli*) and *Staphylococcus aureus* (*S. aureus*) were treated with sub-MICs of reference antibiotics (see Supplementary Table 1). Antibiotics were selected to cover a wide diversity of chemical and pharmacological classes. Another important criterion was the capability of the method to detect weak antibiotic activity. Therefore, assay concentrations were selected to include the MIC and fractions thereof, down to 1/32×MIC, in order to explore the dynamic range of the method. Antibiotic treatment was standardized to the MIC, as the absolute concentration (in this context usually expressed in mg/L) can vary by several orders of magnitude. For example, the MIC of vancomycin (256 mg/L) and ciprofloxacin (0.03 mg/L) for *E. coli* vary by a factor of 8000 (Stock and Wiedemann 1999). In typical compound screenings with a single fixed concentration, the compounds’ efficacy is unknown beforehand. This leads to missed hits in the region of low relative activity. By including the effect of antibiotics at a fraction of the MIC, we aimed to obtain information on drugs that have weak activity and might not be detectable by other phenotypic screening methods.

Bacterial culturing, compound treatment and MALDI-TOF MS were performed in 384-well format. Mass spectral pre-processing was followed by data-dependent feature selection to identify peaks that showed considerable changes in relative intensity upon treatment with antibiotics. Peaks selected for the different models are listed in Supplementary Table 3 (E. coli data) and Supplementary Table 4 (S. aureus data). An exemplary mass spectrum (Figure 1A) and details of two selected peaks for E.coli are depicted in Figure 1B-C, and the corresponding data for S. aureus is provided in Supplementary Figure 1A-C. Using the selected subsets of peaks, quadratic support vector machine classification models (Q-SVM) were trained and internally validated using stratified 10-fold cross validation and stratified 34% hold-out validation. A summary of the evaluated models and their corresponding performance during internal and external validation is listed in Figure 1H. Binary classifiers were trained to identify whether spectra belonged to cell cultures treated with or without an antibiotic. Thus, the total data set for the binary classifiers contained spectra obtained for all seventeen antibiotics at all assayed concentrations (1× to 0.031×MIC in 2-fold dilution series). As an example, the confusion matrix of the 10-fold cross validated binary Q-SVM model of E. coli is given in Figure 1E, providing classification details of 908 mass spectra obtained for all antibiotics at all measured concentrations. In addition, multiclass models were trained with the mode of action as class labels. Antibiotics were grouped to the same classes based on the distinction of their target sites: cell wall synthesis, CWL; protein synthesis, PRT; nucleic acid synthesis processing, DNA; or other mode of action, OTH. The confusion matrix of the 10-fold cross validated mode of action model of E. coli is given in Figure 1G. Details of internal validation of models on S. aureus data are provided in Supplementary Table 5, Supplementary Table 6, Supplementary Table 7 and Supplementary Table 8. Moreover, mass spectra can paint an even more finely grained picture, as it allows for making the distinction between antibiotics of the same class. We show that PhenoMS-ML is able to distinguish between interference in cell wall synthesis caused by vancomycin and the interaction with penicillin-binding proteins by the β-lactams. Within the group of β-lactams, a further discrimination of target profiles is possible, even at a fraction of the MIC (0.125×MIC, see Figure 1F). Similarly, we were able to distinguish (at 0.063×MIC) different target sites on bacterial ribosomes, which are difficult to investigate by biochemical methods, see Supplementary Table 9.

**Figure 1.**
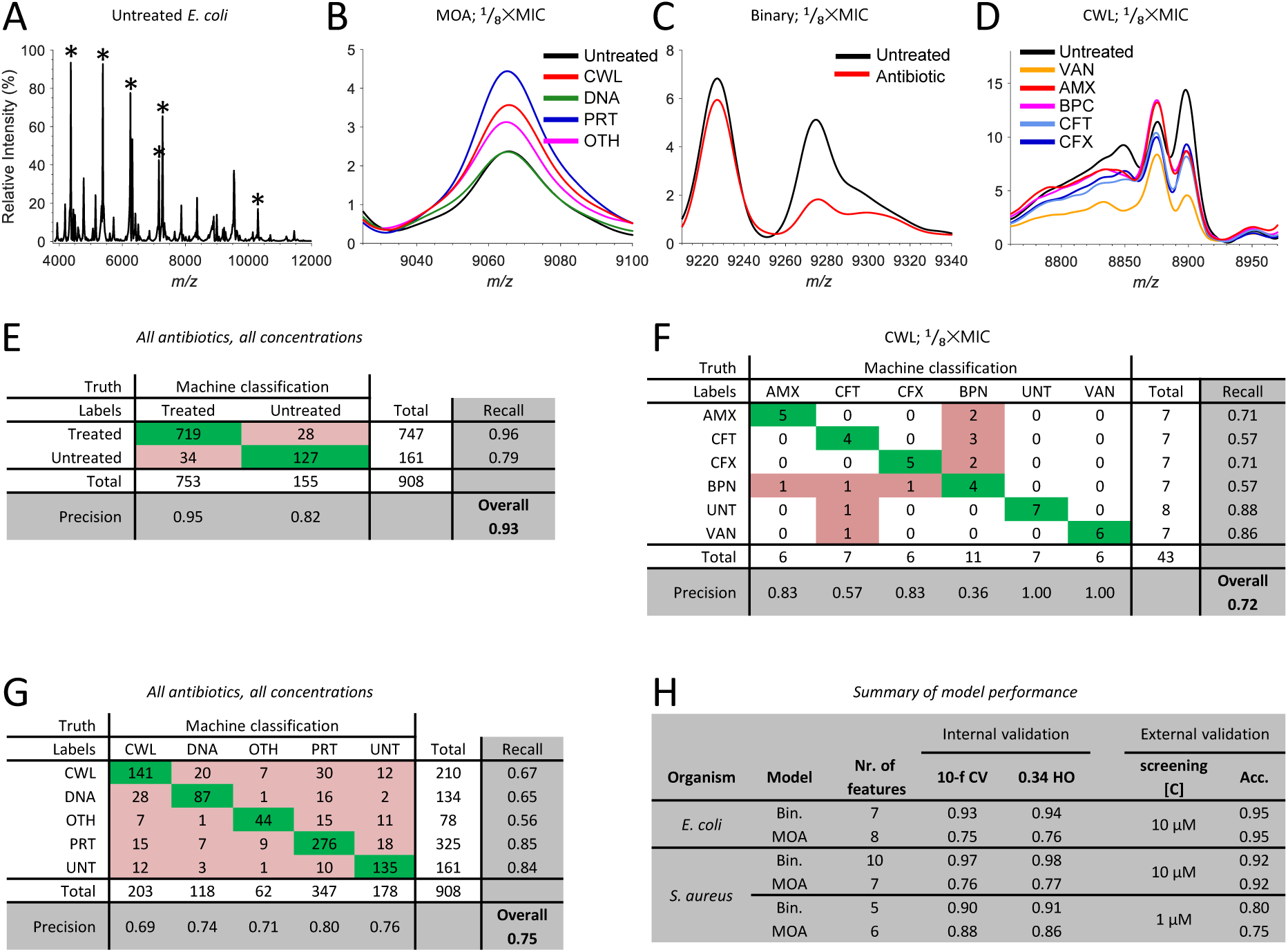
(A) Average mass spectrum of E.coli. Indicated with asterisk (*) are reference peaks used for spectral alignment in the mass spectral pre-processing steps. Details of these high-abundant reference peaks are provided in Supplementary Table 2. (B) Detail of peak at m/z 9065.6, selected for mode of action classification model. Spectra depicted are averaged from the training data set at ⅛×MIC. The relative intensity of this peak in relation to untreated cells (UNT, black) increases upon treatment with antibiotics of the protein synthesis inhibitor (PRT, blue), cell wall synthesis inhibitor (CWL, red) and other antibiotics (OTH, magenta) classes, but not when treated with antibiotics of the nucleic acid synthesis and repair (DNA, green) class. The peak m/z 9065.6 was tentatively identified as acid stress chaperone HdeB (for details see Supplementary Table 3), a protein known to be involved in stress response of E.coli (Kern, Malki et al. 2007). Details of the mode of action classification model with all concentration data is given in G. (C) Detail of peak at m/z 9293.5, selected for the binary classification model. Mass spectra depicted are average mass spectra of all antibiotics in training data set at ⅛×MIC. Relative intensity of the peak at m/z 9293.5 decreases when treated with antibiotics (red) compared to untreated spectra (black), regardless of antibiotic class or concentration. Note that for this subset of spectra at ⅛×MIC, the change of peak intensity is even more pronounced for the peak at m/z 9275.2. However, the data-dependent feature selection did not elect the latter peak for inclusion in modeling when considering all the spectra at all the assayed concentrations. Details of the binary classification model with all concentration data is given in E. (D) Close-up of peaks at m/z 8848.8 and m/z 8897.9, both selected for the antibiotic identity multiclass classification model within the subgroup of cell wall synthesis inhibitors. Depicted is the average mass spectrum of untreated cells (black) and the mass spectra of cells treated with vancomycin (VAN, orange), the β-lactams amoxicillin (AMX, red), benzylpenicillin (BPN, magenta), cefotaxime (CFT, light blue), and cefuroxime (CFX, dark blue). Note the differential responses of the spectral profiles against β-lactams versus vancomycin (m/z 8897.9).Even within the β-lactam group, a differential response can be observed at m/z 8848.8, where cephalosporins cause a decrease and penicillins an increase of relative intensity. Details of the corresponding classification model are given in F. (E) Confusion matrix for the 10-fold cross validated binary Quadratic Support Vector Machine (Q-SVM) model of *E. coli*, representing 908 mass spectra of all assayed antibiotics, at all concentrations. (F) Confusion matrix for the 10-fold cross validated cell wall synthesis inhibitors Q-SVM model of *E. coli*, assayed at ⅛×MIC. Confusion matrix accompanies data depicted in D. (G) Confusion matrix for the 10-fold cross validated mode of action Q-SVM model of *E. coli*, representing 908 mass spectra of all assayed antibiotics, at all concentrations (H) Summary of model performances for both *E. coli* and *S. aureus* during internal and external validation of the binary (Bin.) and mode of action (MOA) models. Listed is the number of features in each model, overall model accuracy using 10-fold cross validation (10-f CV) and 34% hold-out validation (0.34 HO). External validation accuracy (Acc.) of the model was performed using the blind set of drugs of which details given in Table 1. For *S. aureus* models are listed twice as the blind screen (and thus the model construction) was repeated at 1 μM because of poor mass spectral signal quality when screening at 10 μM (see material and methods for details).

Models were externally validated with a blind set of drugs, unknown to the operator of the method. This set of blind drugs included antibiotic and non-antibiotic compounds, to assess both the binary and mode of action classifiers. The binary model of *E. coli* was able to classify 95% of the mass spectra to the correct class. Only the spectrum of cells treated with tiamulin was inadvertently assigned as being untreated by the model. The mode of action model had an overall accuracy of 95% as well. Interestingly, the mode of action model did correctly classify the spectrum from cells treated with tiamulin as being treated with a protein synthesis inhibitor. The mode of action model only inadvertently classified the spectrum from cells treated with nalidixic acid as being treated with a protein synthesis inhibitor. Details of the external validation of models for *E. coli* data are provided in Table 1**Error! Reference source not found.**. Overall accuracy of binary and mode of action models during external validation for S. aureus is comparable to *E. coli.* Details of the external validation of the models for *S. aureus* are provided in Supplementary Table 10. An aspect recognized here is that the predictive power extends beyond the recognition of target sites in the training set. The external validation set also included two probes (tiamulin and fusidic acid) that interfere with target sites (peptidyl transferase unit of the 50S ribosomal subunit and the turnover of elongation factor-G from the ribosome, respectively) not included in model training.

**Table 1.** Details of the predictions made by classification models of *E. coli* during external validation on the blind data set. Indicated in the second column (10 μM/MIC) at which fraction of the MIC the antibiotics were dosed during the screen at 10 μM. Check mark (✓) indicates correct predictions with respect to the expected classification of the model. Details of incorrect predictions are stated in brackets. Overall performance of both models is evaluated using the overall accuracy, indicated at the bottom.

| Nr. of features |  |  | 7 | 8 |
| --- | --- | --- | --- | --- |
| Drug name | 10 μM/MIC | Expected classification | Binary | MOA |
| Brucine | NA | Inactive | ✓ | ✓ |
| Ephedrine | NA | Inactive | ✓ | ✓ |
| Ergotamine | NA | Inactive | ✓ | ✓ |
| Fenbendazole | NA | Inactive | ✓ | ✓ |
| Loperamide | NA | Inactive | ✓ | ✓ |
| Metoprolol | NA | Inactive | ✓ | ✓ |
| Paroxetine | NA | Inactive | ✓ | ✓ |
| Sumatriptan | NA | Inactive | ✓ | ✓ |
| Thalidomide | NA | Inactive | ✓ | ✓ |
| Umifenovir | NA | Inactive | ✓ | ✓ |
| Ampicillin | 0.44<sup>c</sup> | Active/CWL | ✓ | ✓ |
| Azithromycin | 0.94<sup>c</sup> | Active/PRT | ✓ | ✓ |
| Cefuroxime | 0.53<sup>c</sup> | Active/CWL | ✓ | ✓ |
| Chlortetracycline | 1.00<sup>h</sup> | Active/PRT | ✓ | ✓ |
| Fusidic acid | NA<sup>a</sup> | Active/PRT | ✓ | ✓ |
| Nalidixic acid | 0.29<sup>c</sup> | Active/DNA | ✓ | (PRT) |
| Novobiocin | 0.02<sup>d</sup> | Active/DNA | ✓ | ✓ |
| Paromomycin | 1.54<sup>e</sup> | Active/PRT | ✓ | ✓ |
| Tiamulin | 0.62<sup>f</sup> | Active/PRT | (Inactive) | ✓ |
| Trimethoprim | 1.45<sup>c</sup> | Active/DNA | ✓ | ✓ |
| Overall accuracy |  |  | 0.95 | 0.95 |

<sup>a</sup> not active on *E. coli*
<sup>b</sup> not active on *S. aureus*
<sup>c</sup> (EUCAST 2018)
<sup>d</sup> Weakly active on *E. coli* (Sanchez and Watts 1999)
<sup>e</sup>(Zhou, Gregor et al. 2005)
<sup>f</sup> (Xu, Zhang et al. 2009)
<sup>h</sup>(Stanton and Humphrey 2003)

PhenoMS-ML offers a straightforward, high-throughput, label-free, and data-dependent access to highly relevant antibiotic target sites. Additional advantages of the PhenoMS-ML procedure are, contrary to typical MS-based assays, that it does not require tryptic digestion of protein samples, nor does it require solvent and time-consuming liquid chromatography steps prior to sample ionization. The resulting classification models reliably identify specific proteomic signatures induced by interference with the most important target sites of antibiotics, such as cell wall metabolism, ribosomal machinery, and nucleic acid processing, which are difficult to interrogate in biochemical assays on isolated target proteins. Notably, biological responses can frequently be observed at low levels of target interference, which allows the identification of weakly active hits with optimization potential. This opens a perspective for fragment-based drug discovery in a phenotypic setting. As indicated by ongoing studies, PhenoMS-ML can be extended towards eukaryotic systems. The combination of mass spectrometry and machine learning in PhenoMS-ML extends the MALDI-TOF mass spectrometry toolbox towards a phenotypic screening of compounds in wild-type cell cultures in a target and species agnostic manner.

## Acknowledgements

This work was funded by the basic governmental funding of Heidelberg University (Germany). We thank H. Rudy, R. Garg and S. Kämmerer for technical assistance.

## Author contributions

L.N.v.O. and C.D.K. conceived the study. L.N.v.O. performed the experiments and data analysis.

L.N.v.O. and C.D.K. wrote the manuscript.

## Competing interests

The method is subject of a PCT patent application by Heidelberg University, with both L.N.v.O and C.D.K. listed as inventors, filed under reference number PCT/EP2018/079221 (currently under review). The patent application covers all aspects of the method described in this work, along with its applicability towards other organisms.

---

**Supplementary Figure 1.**
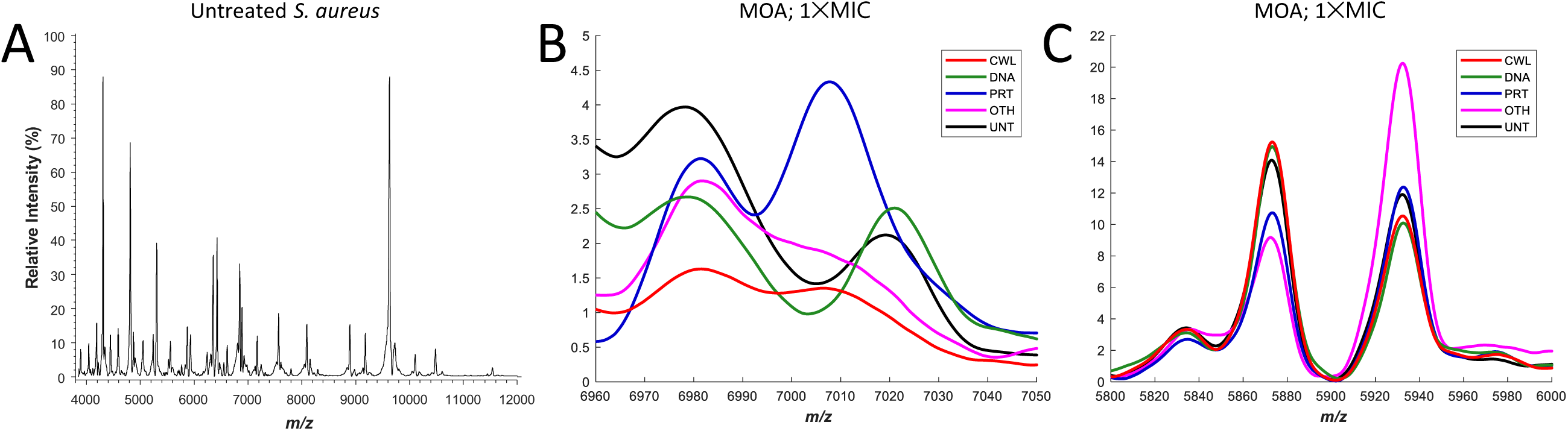
(A) Average mass spectrum of untreated Staphylococcus aureus cell cultures. (B) Enlargement of the m/z 6960-7050 region, with important features at m/z 6978, m/z 7007 and m/z 7020. These three features were selected by the feature selection algorithms for multiple models (see Supplementary Table 4). The depicted mass spectra are average mass spectra of cell cultures treated with 1×MIC of a representative antibiotic of each class: amoxicillin (CWL, red), ciprofloxacin (DNA, green), erythromycin (PRT, blue), nitrofurantoin (OTH, magenta), and untreated (UNT, black) cells. Note especially the peak at m/z 7007, which is only present in spectra of cells treated with antibiotics of PRT class. (C) Detail of peaks at m/z 5873.1 and m/z 5932.5 (tentatively identified as RL33.1 and RL33.2 respectively, see Supplementary Table 4) both selected for the mode of action model of *S. aureus* for the screen at 10 μM. Interestingly, the peak at m/z 5932.5 shows little variation in relative intensity for all antibiotics compared to untreated, except upon treatment with an antibiotic of the class OTH. In that case, the relative intensity of this peak approximately doubles.

**Supplementary Table 1.** List of antibiotics and their respective minimal inhibitory concentrations (MIC, in mg/L) for *S. aureus* and *E. coli*. The accompanying 3-letter abbreviation (Abbr.) for the antibiotic and its general mode of action (MOA) is listed as well.

| Antibiotic | Abbr. | MOA | MIC *E. coli* | MIC *S. aureus* |
| --- | --- | --- | --- | --- |
|  |  |  | ATCC 29522<br>(mg/L) | ATCC 29213<br>(mg/L) |
| Amoxicillin | AMX | CWL | 8 | 2 |
| Benzylpenicillin | PBN | CWL | 32 | 4 |
| Cefotaxime | CFT | CWL | 0.031 | 1 |
| Cefuroxime | CFX | CWL | 8 | 1 |
| Chloramphenicol | CHL | PRT | 8 | 8 |
| Ciprofloxacin | CIP | DNA | 0.004 | 0.25 |
| Clarithromycin | CLR | PRT | 16 | 0.50 |
| Doxycycline | DOX | PRT | 2 | 0.50 |
| Erythromycin | ERT | PRT | 32 | 0.25 |
| Gentamicin | GNT | PRT | 1 | 4 |
| Moxifloxacin | MOX | DNA | 0.064 | 0.008 |
| Neomycin | NEO | PRT | 2 | 8 |
| Rifampicin | RIF | OTH | 16 | 0.008 |
| Tetracycline | TET | PRT | 1 | 1 |
| Trimethoprim | TRM | DNA | 2 | 8 |
| Vancomycin | VAN | CWL | 128 | 2 |
| Nitrofurantoin | NIT | OTH | 16 | 64 |

**Supplementary Table 2.** Reference peaks used for spectra alignment during spectral processing, with their respective protein name and observed and theoretically calculated m/z. RL corresponds to Ribosomal Large subunit (50S) and RS to Ribosomal Small subunit (30S) followed by the respective ribosomal subunit number. Absolute mass accuracy is listed in ppm.

| Name | UniProtKB | Theoretical<br>$m/z$ | Observed<br>$m/z$ | Error<br>(ppm) | Theoretical<br>pI |
| --- | --- | --- | --- | --- | --- |
| RL36 | P0A7Q6 | 4365.3 | 4365.9 | 139 | 10.7 |
| RL34 | P0A7P5 | 5381.4 | 5382.2 | 145 | 13.0 |
| RL33 | P0A7N9 | 6255.4 | 6256.2 | 132 | 10.2 |
| RL32 | P0A7N4 | 6316.2 | 6316.4 | 39 | 11.0 |
| RL35 | P0A7Q1 | 7158.7 | 7159.3 | 74 | 11.8 |
| RL29 | P0A7M6 | 7274.5 | 7275.0 | 75 | 10.0 |
| RS19 | P0A7U3 | 10300.1 | 10300.7 | 59 | 10.5 |

**Supplementary Table 3.** Peaks selected from *E. coli* spectra for the binary and the mode of action (MOA) model. Several peaks that were selected for modelling were identified using the TagIdent tool. Indicated are the theoretical m/z and pI, calculated from the primary amino acid sequence and the corresponding mass accuracy in ppm. Post translational modifications (PTMs) are indicated as well.

| Model | Observed<br>m/z | Theoretical<br>m/z | $\Delta$ Error<br>(ppm) | Name; notes; PTMs | Theoretical<br>pI |
| --- | --- | --- | --- | --- | --- |
| Binary | 4213.9 | - | - | - | - |
|  | 4858.8 | 4859.8 | -202 | Uncharacterized protein YqgB; response to acidic pH | 9.2 |
|  | 7216.1 | 7215.2 | 126 | UPF0253 protein YaeP; uncharacterized protein family | 4.5 |
|  | 7661.0 | - | - | - | - |
|  | 8119.7 | 8119.4 | 35 | Translation initiation factor IF-1; initiator methionine removed | 9.2 |
|  | 8898.3 | - | - | - | - |
|  | 9293.5 | 9293.8 | -32 | Uncharacterized ferredoxin-like protein YfaE | 4.9 |
|  | 12654.5 | 12654.4 | 5 | Ribosome-associated inhibitor A; general response element, , initiator methionine removed | 6.2 |
| MOA | 5097.7 | 5096.8 | 165 | Stationary-phase-induced ribosome-associated protein | 11.0 |
|  | 5411.2 | - | - | - | - |
|  | 6256.2 | 6255.4 | 132 | 50S ribosomal protein L33; initiator methionine removed, N-terminal methylated | 10.2 |
|  | 6280.2 | - | - | - | - |
|  | 6504.2 | - | - | - | - |
|  | 9065.6 | 9066.3 | -71 | Acid stress chaperone HdeB; matured, pos. 30-108 | 4.9 |
|  | 9720.3 | - | - | - | - |

**Supplementary Table 4.**
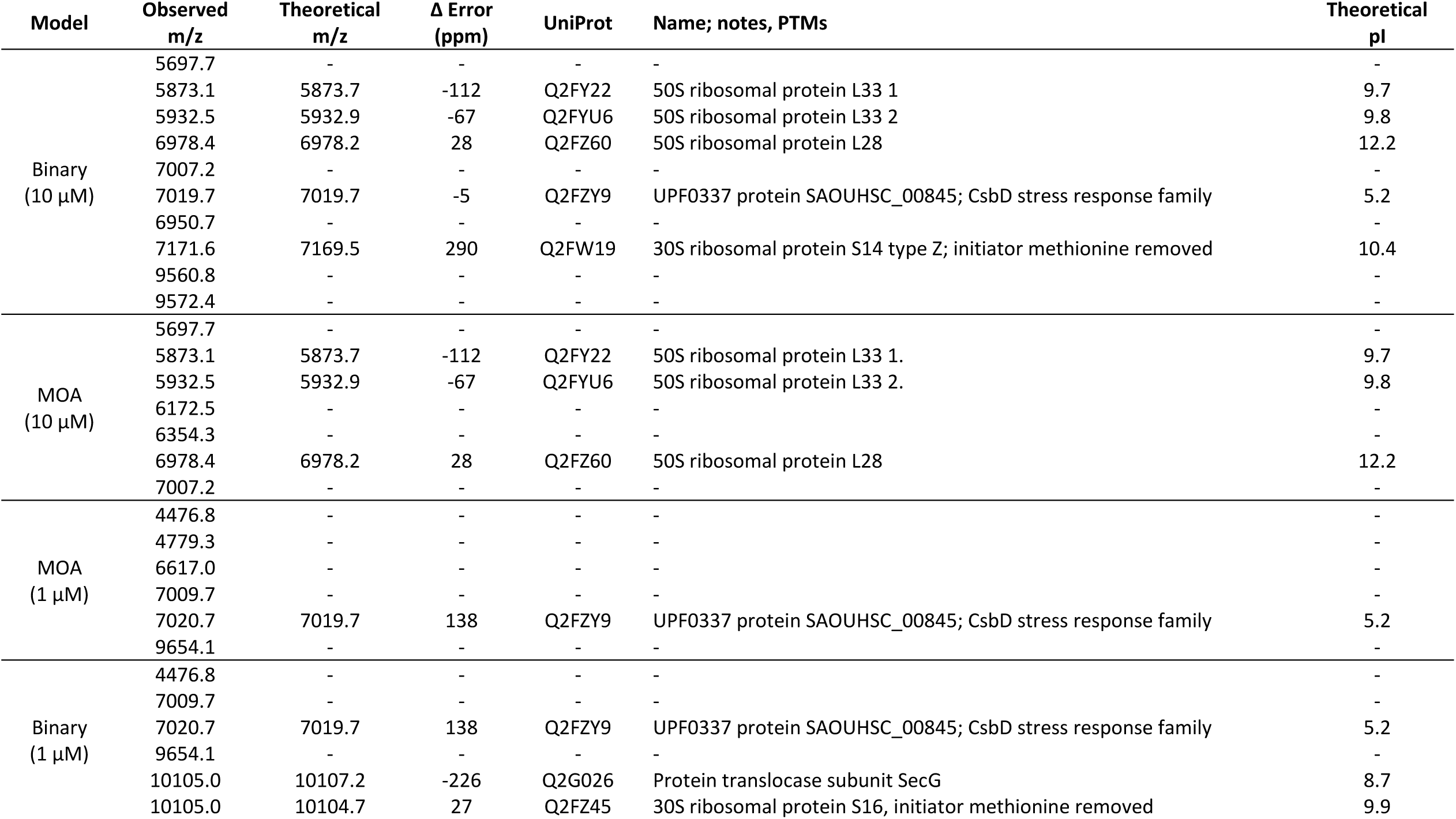
Peaks selected from *S. aureus* spectra for the binary and the mode of action model (MOA) for external validation screens at 10 μM and 1 μM. Several peaks that were selected for modelling were identified using the TagIdent tool. Indicated are the theoretical m/z and pI, calculated from the primary amino acid sequence and the corresponding mass accuracy in ppm. Post translational modifications (PTMs) are indicated as well.

**Supplementary Table 5.** Confusion matrix of the 10-fold cross validation of binary Quadratic Support Vector Machine model of *S. aureus*, representing 860 mass spectra (all antibiotics at all concentrations). This particular model was externally validated with blind drugs screened at 10 μM, of which the details can be found in Supplementary Table 10.

| Truth Labels | Machine classification |  | Total | Recall |
| --- | --- | --- | --- | --- |
|  | Treated | Untreated |  |  |
| Treated | 679 | 15 | 694 | 0.98 |
| Untreated | 7 | 159 | 166 | 0.96 |
| Total | 686 | 174 | 860 |  |
| Precision | 0.99 | 0.91 |  | **Overall 0.97** |

**Supplementary Table 6.** Confusion matrix of the 10-fold cross validation of mode of action Quadratic Support Vector Machine model of *S. aureus*, representing 860 mass spectra (all antibiotics at all concentrations). This particular model was externally validated with blind drugs screened at 10 μM, of which the details can be found in Supplementary Table 10.

| Truth<br>Labels | Machine classification |  |  |  |  | Total | Recall |
| --- | --- | --- | --- | --- | --- | --- | --- |
|  | CWL | DNA | OTH | PRT | UNT |  |  |
| CWL | 160 | 18 | 6 | 22 | 6 | 212 | 0.75 |
| DNA | 18 | 78 | 3 | 32 | 3 | 134 | 0.58 |
| OTH | 2 | 10 | 41 | 18 | 15 | 86 | 0.48 |
| PRT | 13 | 27 | 7 | 215 | 0 | 262 | 0.82 |
| UNT | 5 | 3 | 0 | 1 | 157 | 166 | 0.95 |
| Total | 198 | 136 | 57 | 288 | 181 | 860 |  |
| Precision | 0.81 | 0.57 | 0.72 | 0.75 | 0.87 |  | **Overall**<br>**0.76** |

**Supplementary Table 7.**
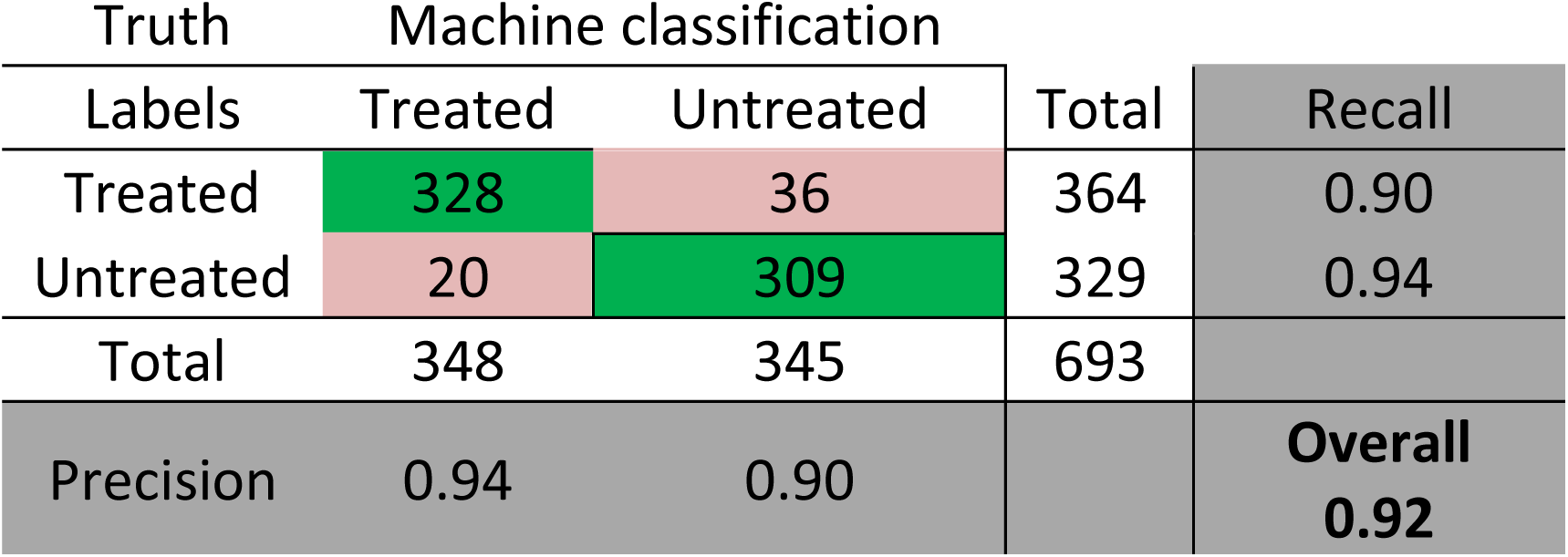
Confusion matrix of the 10-fold cross validation of binary Quadratic Support Vector Machine model of *S. aureus*, representing 693 mass spectra of *S. aureus*(fewer amount of antibiotics included than for screen at 10 μM, at 1×, 0.5×, 0.25× and 0.125×MIC, see material and methods for details). This particular model was externally validated with blind drugs screened at 1 μM, of which the details can be found in Supplementary Table 10.

| Truth<br>Labels | Machine classification |  | Total | Recall |
| --- | --- | --- | --- | --- |
|  | Treated | Untreated |  |  |
| Treated | 328 | 36 | 364 | 0.90 |
| Untreated | 20 | 309 | 329 | 0.94 |
| Total | 348 | 345 | 693 |  |
| Precision | 0.94 | 0.90 |  | **Overall**<br>**0.92** |

**Supplementary Table 8.** Confusion matrix of the 10-fold cross validation of mode of action Quadratic Support Vector Machine model of *S. aureus*, representing 693 mass spectra of *S. aureus* (fewer amount of antibiotics included than for screen at 10 μM, at 1×, 0.5×, 0.25× and 0.125×MIC, see material and methods for details). This particular model was externally validated with blind drugs screened at 1 μM, of which the details can be found in Supplementary Table 10.

| Truth | Machine classification |  |  |  |  | Total | Recall |
| --- | --- | --- | --- | --- | --- | --- | --- |
| Labels | CWL | DNA | OTH | PRT | UNT |  |  |
| CWL | 97 | 1 | 0 | 5 | 7 | 110 | 0.88 |
| DNA | 1 | 43 | 2 | 7 | 3 | 56 | 0.77 |
| OTH | 0 | 1 | 13 | 3 | 5 | 22 | 0.59 |
| PRT | 1 | 9 | 2 | 143 | 21 | 176 | 0.81 |
| UNT | 3 | 4 | 1 | 8 | 313 | 329 | 0.95 |
| Total | 102 | 58 | 18 | 166 | 349 | 693 |  |
| Precision | 0.95 | 0.74 | 0.72 | 0.86 | 0.90 |  | Overall<br>0.88 |

**Supplementary Table 9.** Confusion matrix for the 10-fold cross validated antibiotic identity Quadratic Support Vector Machine model of *E. coli*, representing 63 mass spectra of cells treated with a variety of protein synthesis inhibitors (chloramphenicol; CHL, clarithromycin; CLR, doxycycline; DOX, erythromycin; ERY, gentamycin; GNT, neomycin; NEO, tetracycline; TET and untreated cells’ mass spectra; UNT) at 0.063×MIC. Note the slight confusion of the model between both aminoglycosides (GNT and NEO) and between tetracyclines (TET and DOX). At this relatively low concentration, the effect of clarithromycin (CLR) becomes more difficult to distinguish from spectra from untreated cells, contributing to a relatively low precision of the class (UNT).

| Truth |  | Machine classification |  |  |  |  |  |  |  |  |
| --- | --- | --- | --- | --- | --- | --- | --- | --- | --- | --- |
| Labels | CHL | CLR | DOX | ERY | GNT | NEO | TET | UNT | Total | Recall |
| CHL | 6 | 0 | 1 | 0 | 0 | 0 | 0 | 0 | 7 | 0.86 |
| CLR | 0 | 5 | 0 | 0 | 0 | 0 | 0 | 3 | 8 | 0.63 |
| DOX | 0 | 0 | 5 | 1 | 0 | 0 | 1 | 1 | 8 | 0.63 |
| ERY | 0 | 0 | 0 | 8 | 0 | 0 | 0 | 0 | 8 | 1.00 |
| GNT | 0 | 0 | 0 | 0 | 7 | 1 | 0 | 0 | 8 | 0.88 |
| NEO | 0 | 0 | 0 | 0 | 1 | 6 | 1 | 0 | 8 | 0.75 |
| TET | 0 | 0 | 0 | 0 | 0 | 0 | 8 | 0 | 8 | 1.00 |
| UNT | 0 | 1 | 0 | 0 | 1 | 0 | 0 | 6 | 8 | 0.75 |
| Total | 6 | 6 | 6 | 9 | 9 | 7 | 10 | 10 | 63 |  |
| Precision | 1.00 | 0.83 | 0.83 | 0.89 | 0.78 | 0.86 | 0.80 | 0.60 |  | Overall<br>0.81 |

**Supplementary Table 10.**
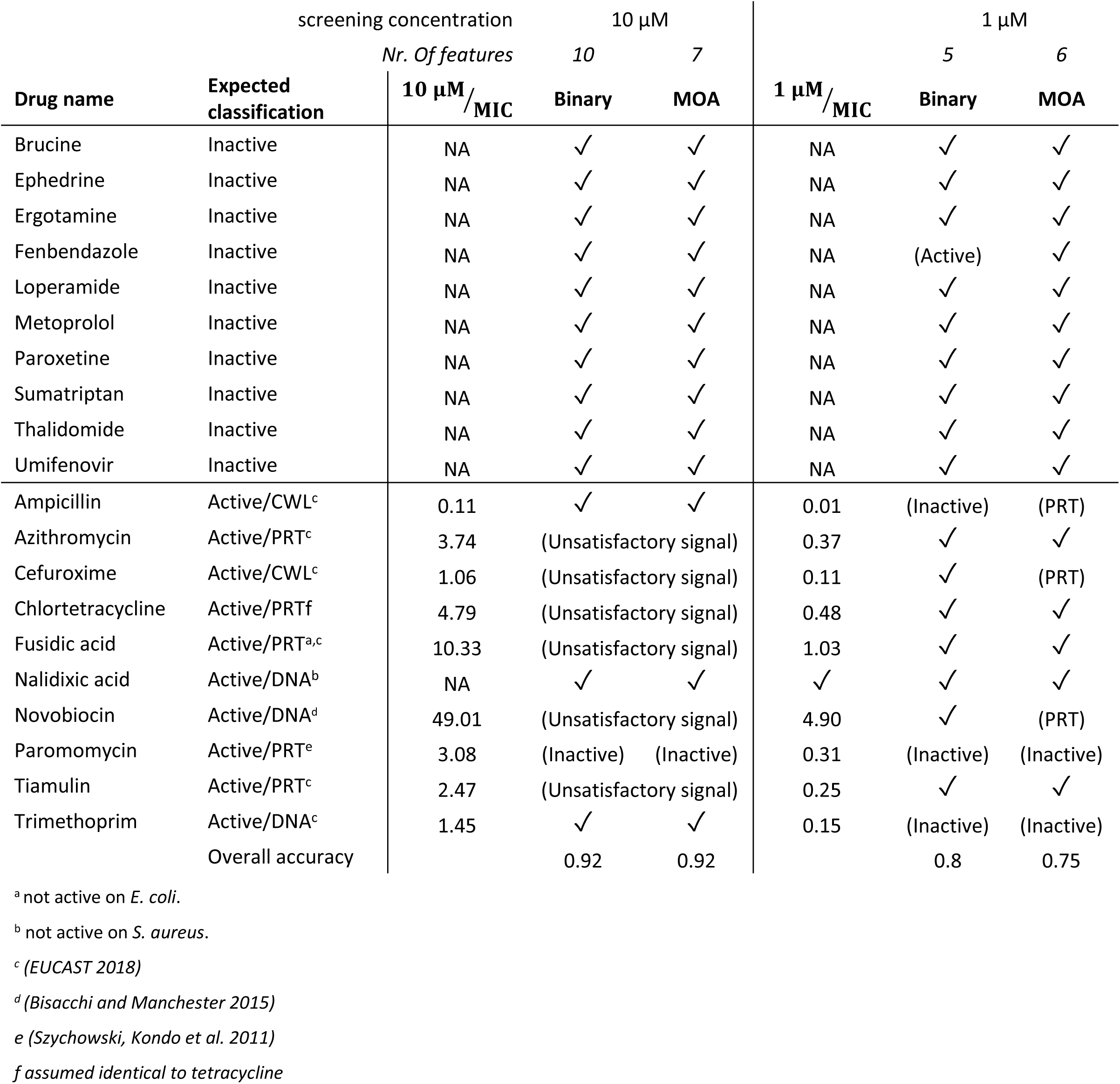
External validation on *S. aureus*. Details of the predictions made by classification models of *S. aureus* during external validation on the blind data set. Check mark (✓) indicates correct predictions, details of incorrect predictions are stated in brackets. At the screening concentration of 10 μM, several spectra were removed from the dataset due to unsatisfactory signal quality, as indicated in the table. This was the case for several antibiotics which were dosed >>MIC. The screening of *S. aureus* was therefore repeated at 1 μM. Overall performance of the models is evaluated using the overall accuracy (indicated at the bottom).

## Material and methods

### Medium and antibiotics

All experiments were performed using cation-adjusted Mueller-Hinton medium (MH medium; Sigma-Aldrich, Munich, Germany) prepared according to the manufacturers’ guidelines. Antibiotics were selected to cover a diverse range of modes of action, listed in Supplementary Table 1. The following antibiotics were dissolved in water: benzylpenicillin (BPN), cefotaxime (CFT), cefuroxime (CFX), moxifloxacin (MOX), and vancomycin (VAN). The following antibiotics were dissolved in dimethyl sulfoxide (DMSO) and water (50 v/v%): amoxicillin (AMX), ciprofloxacin (CIP), erythromycin (ERY), gentamicin (GNT), neomycin (NEO), tetracycline (TET), trimethoprim (TRM), nitrofurantoin (NIT), and rifampicin (RIF). The following antibiotics were dissolved in DMSO: chloramphenicol (CHL), clarithromycin (CLR), and doxycycline (DOX). Antibiotics were dissolved to a final concentration of 1280 mg/L and filtered using a cellulose acetate membrane (0.2 µm pore size, GE Healthcare Life Science, Freiburg, Germany) to ensure sterility. Stock solutions were stored at 4° Celsius. Prior to use, antibiotic stock solutions were diluted in sterile cation-adjusted MH medium.

### MIC determination

The MICs of selected antibiotics were determined in accordance with the CLSI (CLSI 2013) and EUCAST (EUCAST 2016) guidelines for antimicrobial susceptibility testing, as described in detail by Wiegand and coworkers (Wiegand, Hilpert et al. 2008).The MIC was determined for the Gram-negative *Escherichia coli* strain (DSMZ 1103, equivalent to ATCC 25922) and the Gram-positive *Staphylococcus aureus* (DSMZ 2569, equivalent to ATCC 29213), obtained from the DSMZ (Deutsche Sammlung von Mikroorganismen und Zellkulturen; German collection of microorganisms and cell cultures).

### Bacterial cell culture synchronization

The replication and division cycles of the bacteria were synchronized. *E. coli* cells were grown in 50 mL tubes for approximately eight hours in MH medium in a Minitron incubator (Infors AG, Bottmingen, Switzerland) at 120 rotations per minute (rpm) with 25 mm shaking throw at 37° C, after which cells were centrifuged at 2000×g for 10 minutes (Rotina 420R, Hettich Lab Technology, Tuttlingen, Germany). Residual medium was decanted to waste and the cell pellet was resuspended in sterile DPBS (Dulbecco’s phosphate buffered saline, Sigma-Aldrich, Munich, Germany). Cell cultures were starved in this nutrient limited environment (120 rpm; at 37° C) overnight for approximately 16 hours. After starvation, cells were centrifuged for 10 minutes at 2000×g. Supernatant was decanted to waste and cells were resupplied with fresh MH medium and diluted to McFarland standard of 1.0. Cells were allowed to adapt to the nutrient rich medium for at least one division cycle (approximately 70 minutes in the case of *E. coli*; approximately 90 minutes in the case of *S. aureus*) to a McFarland of 2.0 before addition to the antibiotics in the 384-well plate at a final cell density with McFarland 1.0, corresponding to 1×10^8^ colony forming units per mL (CFU/mL).

### Antibiotic treatment

The concentrations at which experiments were performed are denoted as a fraction of the MIC in the following manner throughout the remainder of this work: for example, ⅛×MIC for an experiment performed at 1/8th of the MIC value (0.125×MIC). Cells were exposed to 1×, 0.5×, 0.25×, 0.125×, 0.063×, and 0.031×MIC, unless indicated otherwise. Eight biological replicate cell cultures per concentration were prepared, to yield eight replicate mass spectra per assayed condition. Exposure of cells to antibiotics was performed in clear polystyrene 384-well plates (flat-bottom; Greiner Bio-One GmbH, Frickenhausen, Germany). Concentrations of each antibiotic (2-fold dilution series in cation-adjusted MH) were made to ensure that the highest final assay concentration was 1×MIC of that antibiotic. First, 50 µL of antibiotic stock (2×MIC) solution were added to each well. Subsequently an inoculum of 50 µL with 2×10^8^ CFU/mL to the plates using a multichannel pipette to ensure final cell density of 1×10^8^ CFU/mL. Plates were sealed using sealing film (SealPlate® film, Excel Scientific Inc, Victorville, CA, USA) and placed in a preheated microplate incubator (Thermo Scientific iEMS Incubator/Shaker, ThermoFisher Scientific, Waltham, MA, USA) at 37° C and shaken at 1150 rpm for 2 hours.

### Sample preparation

After incubation, 384-well plates were centrifuged (Rotina 420R, Hettich Lab Technology, Tuttlingen, Germany) equipped with a swinging bucket rotor at 2000×g for 10 minutes. Supernatant was discarded and cell pellets were washed with 100 µL 35% ethanol (v/v%) and incubated in the microplate incubator for 5 minutes at 1150 rpm. Cell debris was centrifuged again and washed a second time with 100 μL of 35% ethanol. After removal of 90 μL the supernatant, cells were resuspended in the remaining 10 µL 35% ethanol, sealed and stored at 4 °C. Prior to MALDI-TOF MS analysis, bacterial cell pellets were resuspended in the plate by shaking in the microplate incubator for 5 minutes at 1150 rpm. Cell suspension was mixed 1:1 with freshly prepared α-cyano-4-hydroxycinnamic acid (CHCA; 10 mg/mL in 50.0% acetonitrile, 47.5% H2O, and 2.5% trifluoroacetic acid) and approximately 1 µL was spotted on a MALDI target plate (MSP 96 polished steel BC microScout target, Bruker Daltonics, Bremen, Germany). Samples were air-dried at room temperature.

### MALDI-TOF settings

Target plates were positioned in the mass spectrometer (MALDI-TOF microflex LT, Bruker Daltonics, Bremen, Germany) fitted with a nitrogen laser (337 nm, set to 60 Hz). Spectra were acquired in linear mode with a mass range of *m/z* 2,000-15,000 using AutoXecute runs of the FlexControl software (Version 3.3, Build 108.2, Bruker Daltonics). The laser was set to fire 100 shots at 80% power per location (attenuator set to 20-30%), while moving in a small spiral raster over 7 locations per sample spot to assure appropriate signal intensity. The sum of 700 shots yielded spectra with ion intensities in the order of 10^4^-10^5^ ion counts for the most abundant ions. Sample rate was set to 1.00 GS/s; detector gain was set to 3.7×; electronic gain was set to 200 mV and Realtime Smooth was disabled. Default delayed ion extraction was fixed at 140 ns. Calibration of the instrument was regularly evaluated using Brukers ‘Protein Calibration Mix I’ and, if necessary, adjusted accordingly.

### Spectral pre-processing

Using Bruker’s FlexAnalysis software, the collected raw spectra were exported to a *.txt file in ASCII format. Subsequently, the spectra were imported in MATLAB (R2018a; The MathWorks Inc., Natick, USA) installed on a desktop PC (i5-4690 CPU @3.50GHz equipped with 16 GB RAM and a 64-bit Windows 7 Professional operating system) and pre-processed as follows. First, spectra were resampled (MATLAB function *msresample*) in order to obtain a homogenous mass/charge (*m/z*) vector for each sample in the range of *m/z* 3850-15000. The baseline of each individual spectrum was estimated and subtracted using a sliding window filter (MATLAB function *msbackadj*). Noise was reduced using locally weighted scatter plot smoothing regression method (commonly referred to as LOWESS filter; MATLAB function *mslowess*). Spectra were normalized to their total ion current (TIC; MATLAB function *msnorm*) and rescaled such that the highest peak in each mass spectrum had a relative intensity of 100%.

### Spectral quality control

The TIC value was used as a measure for spectral quality. This eliminates the requirement to visually inspect each spectrum, which is a laborious and subjective task. Instead, the TIC allows for an objective verdict about the signal quality of the mass spectrum. Based on the TIC values of the whole dataset, the data was grouped into quartiles and the interquartile range (IQR) of the TIC was calculated. To determine outliers spectra from the bulk TIC data, the upper fence (UF) and the lower fence (LF) were computed using Equation 1 and Equation 2, as described previously by Tukey and coworkers (Tukey 1977, Hoaglin, Iglewicz et al. 1986).

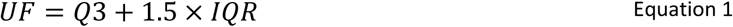

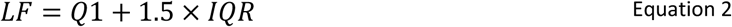

In Equation 1 and Equation 2, Q3 represents the third quartile (75th percentile) and Q1 the first quartile (25th percentile) of the TIC values. Spectra with TIC values above the upper fence or below the lower fence were considered outliers and removed from the dataset.

In addition, an outlier filter was added that removes any spectrum whose intensity was higher than the upper fence based on the intensity of the mass spectrum at *m/z* 12500 (where no peak was observed).

Therefore, the relative intensity at this *m/z* provides an easy way of removing spectra with poor signal quality. As a threshold, spectra with relative intensity above the third quartile plus two times the interquartile range at *m/z* 12500 (where no peak is expected) were removed. In practice, this threshold meant that all spectra with intensity roughly above 1-1.5% at *m/z* 12500 were removed.

### Peak alignment and peak detection

Each mass spectrum was aligned towards known, conserved, and high intensity peaks (MATLAB function *msalign*). The majority of the proteins that can be observed in a typical *E. coli* mass spectrum are large and small ribosome-associated proteins (RL and RS) (Arnold and Reilly 1999). By aligning spectra during the initial processing step towards several of these highly intense and consistently observed peaks, errors in peak location are reduced. In the case of mass spectra of *E. coli*, the peaks used for alignment were observed at the following *m/z* values (protein name; UniProt accession number in parenthesis, post translational modification if applicable): 4365.333 (RL36; P0A7Q6), 5381.396 (RL34; P0A7P5), 6255.416 (RL33; P0A7N9 initiator methionine removed, methylated), 6316.197 (RL32; P0A7N4, initiator methionine removed), 7158.746 (RL35; P0A7Q1, initiator methionine removed), 7274.456 (RL29; P0A7M6) and *m/z* 10300.100 (RS19; P0A7U3, initiator methionine removed). Peaks were putatively identified by searching the UniProt database (release 2018_07) of reference proteome up000000625 of *Escherichia coli* strain K12 (Taxonomy identifier 83333) using the TagIdent tool (Gasteiger, Hoogland et al. 2005). Subsequently, average masses and theoretical pI’s of proteins were calculated using the primary sequence data and the Fragment Ion Calculator (Proteomics Toolkit, Institute for Systems Biology, available at http://db.systemsbiology.net:8080/proteomicsToolkit/FragIonServlet.html).

For *S. aureus*, peak identities were found in the UniProt database using the reference proteome up000008816 of *Staphylococcus aureus* strain NCTC 8325. The peaks of *S. aureus* used for alignment were observed at the following *m/z* values (protein name; UniProt accession number in parenthesis, post translational modification if applicable): *m/z* 4306.36 (RL36; Q2FW29), 5303.35 (RL34; Q2FUQ0, initiator methionine removed), 5873.74 (RL33; Q2FY22), 6354.35 (RL32; Q2FZF1, initiator methionine removed), 6554.68 (RL30; P0A0G2), and *m/z* 9627.02 (DNA-binding protein HU; Q5HFV0). Theoretical average masses were calculated as described for *E. coli*.

A peak detection algorithm based on the undecimated discrete wavelet transform was applied on the average spectrum of replicate experiments to identify centroid peak locations (Coombes, Tsavachidis et al. 2005, Morris, Coombes et al. 2005) (MATLAB function *mspeaks*). Subsequently, peak binning was performed to obtain a common *m/z* vector to describe the peaks observed in the spectra. This yielded a common *m/z* vector containing approximately 170 peaks in the *m/z* 3850-15000 Da region in the case of *E. coli*. A comparable number of peaks is observed for mass spectra of *S. aureus* (∼130 peaks).

Computational time was approximately 2.35 seconds per spectrum, from importing the raw *.txt until peak detection using the mentioned computer and settings.

### Feature selection

Not all peaks in the mass spectra contain sufficient discriminatory information for model construction. Peaks may be removed from the dataset, as some peaks might cause overcomplicating and overfitting (poor generalization) of the models. Therefore, two types of feature selection algorithms have been applied in order to remove noisy and redundant peaks: (1) a random forest (RF) of decision trees and (2) sequential (forward; SFS and backward; SBS) feature selection. Features selected by two or all three of the applied feature selection methods (RF, SFS, and SBS) were considered for final model building.

Firstly, relative classification power of the peaks was evaluated using a random forest of decision trees, a so-called ‘embedded’ feature selection method (Breiman 2001). A bootstrap aggregated (‘bagged’) random forest of 1000 decision trees was grown to evaluate the feature importance (MATLAB function *TreeBagger*). The amount of 1000 trees gives a good estimation of the feature importance considering the data size and complexity (Oshiro, Perez et al. 2012). By evaluating the out-of-bag error, the relative importance of each peak regarding its impact on classification performance was evaluated. As a threshold, features with a relative feature importance higher than the mean importance plus one and a half standard deviation of the mean feature importance were considered for incorporation in the models. This evaluation of feature importance was performed for two different scenarios with different class labelling: (a) by using binary labelling of the data: spectra were labelled either as ‘treated’ or ‘untreated’ with an antibiotic, regardless of antibiotic mode of action or antibiotic concentration. The second labelling (b) was done according to antibiotic mode of action: ‘CWL’ for cell wall synthesis inhibitors, ‘PRT’ for protein translation inhibitors, ‘DNA’ for antibiotics interfering with DNA synthesis and maintenance, ‘OTH’ for other mode of action or ‘No activity’ for untreated cells; regardless of antibiotic concentration.

Subsequently, sequential feature selection (a ‘wrapper’ method) was used to select a subset of peaks that best classifies the data. Features considered for sequential feature selection were features that had a relative feature importance higher than the mean feature importance minus one standard deviation as determined by the RF. This was done in order to reduce calculation time, as sequential feature selection is a computationally expensive method.

During sequential feature selection, a subset of features was selected that best classified the data until there was no improvement in classification accuracy. This was done by creating an initial empty feature subset and subsequently adding more features (MATLAB function *sequentialfs*). Additionally, SBS was performed, where initially all features (that is: only the features with a relative feature importance higher than the mean feature importance minus one standard deviation as determined by the RF) were considered. In that case, features were removed from the initial subset, until accuracy no longer improved. For each new candidate feature subset (after adding or removal of a feature), a stratified 10-fold cross validation was performed. SFS selection was performed 100 times. Features were selected based on the mean amount of times they were selected (out of these 100 times) plus one and a half standard deviation of the amount of times they were selected.

SBS selection was also performed 100 times using stratified 10-fold cross validation. As a threshold, features that were selected more than the mean amount of times they were selected (out of the 100 times) plus one standard deviation of the amount of times they were selected, were considered for modelling. If either the selection threshold for SFS or SBS was >100, which would result in no features selected, a threshold of >99 was taken.

Within the subgroup of cell wall synthesis inhibitors, at ⅛×MIC, features were selected in order to further discriminate between the β-lactams and vancomycin. Due to the relatively small amount of spectra in this particular subgroup, features were only evaluated using a random forest of decision trees. The subgroup of protein synthesis inhibitors was also investigated at a fraction of the MIC (0.063×MIC) and only evaluated using a random forest of decision trees.

### Model building and internal validation

Using the selected features and corresponding class labels (either the drug compound had ‘activity’ or ‘no activity’, or the mode of action, or the compound identity, as listed in Supplementary Table 1), models were constructed under MATLAB’s default settings in the *classificationLearner* application. It was found that quadratic Support Vector Machine-based (Q-SVM) classifying models performed among the best on our data sets. Therefore, in this work only Q-SVM models are discussed. The models were internally validated using a stratified 10-fold cross-validation and stratified 34% hold-out validation.

### Model evaluation

Model performance was evaluated with the overall accuracy, a number between 0 and 1, indicating the fraction of spectra classified correctly (see Equation 3). In addition, for each class in the models, the recall and precision for each class are given, calculated according to Equation 4 and Equation 5 respectively.

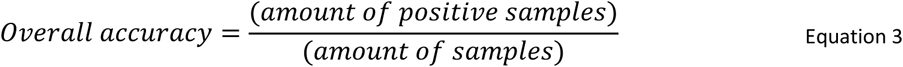

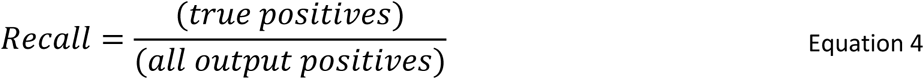

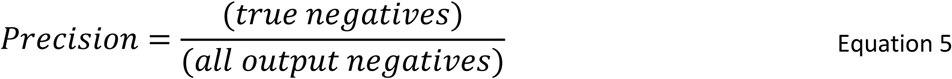

### External validation

The trained models were externally validated by classifying the mode of action on novel data, which was explicitly not included in the model training phase. External validation was performed with a blind set of twenty compounds. These compounds were provided without any further information about their (mode of) activity, only that there were antibiotics and inactive compounds among them. These compounds were subjected to the PhenoMS-ML method, at a fixed concentration of 10 μM, a typical concentration in HTS campaigns. For the validation, two models were built for each bacterial strain. One using a binary classifier, returning only whether the spectra belonged to cells treated with an antibiotic (outcome ‘yes’) or is untreated (outcome ‘no activity’), and a second model that was built used the mode of action of the antibiotics as class labels (as listed in Supplementary Table 1).

In the case of *S. aureus*, treatment of cells with some of the compounds yielded spectra that were deemed of insufficient quality and therefore no classification could be performed. In these instances, it was assumed that the spectra were of insufficient quality due to the fact that the cells were treated with such copious amounts of antibiotic that insufficient cells had grown to generate a signal. These compounds were screened again, but at 1 μM screening concentration instead of 10 μM. For logistic reasons, the training set was reduced slightly: ciprofloxacin, vancomycin, trimethoprim, tetracycline, and nitrofurantoin were excluded for model training.

## Captions

Supplementary Table 1. List of antibiotics and their respective minimal inhibitory concentrations (MIC, in mg/L) for S. aureus and E. coli. The accompanying 3-letter abbreviation (Abbr.) for the antibiotic and its general mode of action (MOA) is listed as well.

